# Evolution of rhodopsin in flatfishes (Pleuronectiformes) is associated with depth and migratory behaviour

**DOI:** 10.1101/2023.11.30.569440

**Authors:** Esme Macpherson, Frances E Hauser, Alexander Van Nynatten, Belinda SW Chang, Nathan R Lovejoy

## Abstract

Visual signals are involved in many fitness-related tasks and are therefore essential for survival in many species. Aquatic organisms are ideal systems to study visual evolution, as the high diversity of spectral properties in aquatic environments generates great potential for adaptation to different light conditions. Flatfishes are an economically important group, with over 800 described species distributed globally, including halibut, flounder, sole and turbot. The diversity of flatfish species and wide array of environments they occupy provides an excellent opportunity to understand how this variation translates to molecular adaptation of vision genes. Using models of molecular evolution, we investigated how the light environments inhabited by different flatfish lineages have shaped evolution in the rhodopsin gene, which is responsible for mediating dim-light visual transduction. We found strong evidence for positive selection in rhodopsin, and this was correlated with both migratory behaviour and several fundamental aspects of habitat, including depth and freshwater/marine evolutionary transitions. We also identified several mutations that likely affect the wavelength of peak absorbance of rhodopsin, and outline how these shifts in absorbance correlate with response to the light spectrum present in different habitats. This is the first study of rhodopsin evolution in flatfishes that considers their extensive diversity, and our results highlight how ecologically-driven molecular adaptation has occurred across this group in response to transitions to novel light environments.

## 1. INTRODUCTION

### 1.1 Visual transduction and aquatic light environments

The visual system allows for extremely rapid detection of environmental stimuli. Because visual signals are involved in many fundamental tasks such as avoiding predators, navigation, communication, foraging, and mating, optimizing the molecular processes involved in visual phototransduction is of great importance to survival and fitness of many organisms (Dungan et al. 2016, Hauser and Chang 2017). The diversity of visual system modifications across the animal kingdom is of longstanding interest to evolutionary biologists, and provides an opportunity to study how natural selection tunes sensory systems to adapt to different ecological conditions. An integral part of this system are the visual pigments: light-sensitive G protein-coupled receptor opsin proteins bound to a light-sensitive retinal chromophore (Hauser and Chang 2017, Yokoyama et al. 1995). Visual pigments are expressed in photoreceptor cells of the retina, and function by absorbing incoming photons of certain wavelengths. The photons trigger a conformational change in the photoreceptor protein, which initiates the visual transduction cascade and ultimately results in a neural signal that light has been detected (Yokoyama et al. 1995). Because opsin proteins act at the interface between the organism and its environment, studying opsin evolution allows examination of the environmental influence on fitness-related molecular adaptation (Hauser and Chang 2017, Hauser et al. 2021).

Aquatic organisms provide an ideal basis to study adaptation in the visual system to different environments, because the unique physical properties of water impose strong selection pressures on the visual system (Hunt et al. 1996, Hauser and Chang 2017). In aquatic systems, the physical properties of water cause considerable differences in the amount and colour of available light at different depths. Most notably, total light decreases steadily with depth to approximately 1000m below the surface, beyond which is near total darkness (Hunt et al. 1996, Luk et al. 2016, Xia et al. 2021). Other changes occur across the visible spectrum: in shallow waters there is a broader spectrum of light available because violet & red-green wavelengths attenuate quickly with depth (Van Nynatten et al. 2021. Additionally, other environmental properties can affect aquatic ambient light environments, such as suspended sediments that increase turbidity (Costa et al. 2013). Shallow coastal, estuarine, and fresh waters can exhibit high degrees of scattering and absorption of light caused by dissolved or suspended particles, producing highly varied photic environments (Costa et al. 2013, Hill et al. 2019,).

### 1.2 Ecology and distribution of flatfishes

Flatfishes (Pleuronectiformes) are a species-rich teleost order, distributed globally in diverse habitats ranging from tidal zones to deepwater benthic communities, and spanning tropical to arctic regions (Gibson 2008, Campbell et al. 2019). Several species are freshwater endemics. Other species migrate between freshwater and marine habitats (anadromy or catadromy), or migrate into the water column in the open ocean (oceanodromy) (Gibson 2008, Bitencourt et al. 2023). Of the more than 800 recognized flatfish species, many are economically important to global fisheries (e.g. halibut, flounder, sole, turbot) (Gibson 2008, Atta et al. 2021). Flatfishes possess a specialized asymmetrical and laterally compressed body plan related to their benthic lifestyle (Gibson 2008, Atta et al. 2021). Larval flatfish begin life upright, but during development undergo a 90° rotation in the body plan, where one eye migrates to the other side of the head and the organism begins swimming on one side (Gibson 2008). Previous analyses of individual flatfish lineages have demonstrated varied modifications in the visual system to benthic life (Iwanicki et al. 2017, Savelli et al. 2018, Frau et al. 2020, Nag et al. 2022), but to date there have been no investigations of opsin evolution across the vast array of flatfishes. The diversity of flatfishes, their global distribution, and the wide variety of environmental conditions they inhabit provides an excellent opportunity to understand how environmental variation translates to molecular adaptation.

### 1.3 Visual opsins

Teleost fishes have complex visual systems, with five classes of visual opsin proteins including rhodopsin (encoded by the RH1 gene) which is involved in scotopic (dim-light) vision, and four cone opsins mediating photopic (colour) vision ranging in maximum absorbance (λ_max_) from ultraviolet to red (Lin et al. 2017, Hauser et al. 2021). Adaptive changes in the number, expression, function, and absorbance wavelength of visual pigments allow different species and life stages to function more efficiently in their ambient light environments (Hauser and Chang 2017). Aquatic lineages that have undergone habitat transitions often show signs of adaptation in the visual system to the novel environmental conditions (Hunt et al. 1996, Hauser et al. 2021, Xia et al. 2021, Dungan and Chang 2022). These include amino acid substitutions in the opsin protein that alter the wavelength of peak absorbance in visual pigments, known as ‘spectral tuning’ mutations (Hauser and Chang 2017, Lin et al. 2017). Certain substitutions at spectral tuning sites are highly suggestive of adaptive evolution in the visual system; examples include blue-shifting mutations in deep-water species (Hunt et al. 1996, Dungan and Chang 2017, Dungan and Chang 2022, Ricci et al. 2022). Other mutations known to have an affect on the opsin’s function can also be signs of ecological adaptation, such as sites that alter the kinetics of opsin activation (i.e., how quickly the opsin may recover following activation; Luk et al. 2021, Xia et al. 2021).

Previous research on flatfish visual opsins has focused on ontogenetic changes in opsin expression during eye migration in larval development (Frau et al. 2020, Helvik et al. 2001, Frau et al. 2022). Some studies have determined that flatfishes do not express rod cells or RH1 in the eyes until post-metamorphosis, although these studies have been limited to individual species (Pankhurst and Butler 1996, Helvik et al. 2001) and are not reflective of all flatfishes (Frau et al. 2020). There is also considerable diversity in the number and spectral sensitivity of cone opsins; evolution in flatfish cone opsins is often associated with adaptation to the variable light environments experienced between pelagic larval and benthic adult stages (Bolstad and Novales Flamarique 2022, Zhang et al. 2022). However, to date, there have no been no comprehensive investigations of lineage-wide patterns of evolution in flatfish vision genes. The dim-light vision gene rhodopsin is particularly well-suited to this type of study, because of its documented importance for adaptation to different habitats and ecology (Hunt et al. 1996, Hauser and Chang 2017, Hauser et al. 2021) and the availability of sequences for a broad range of taxa.

Here, we investigated the molecular evolution of the RH1 gene across the entire flatfish order, incorporating 173 species from all 16 flatfish families. We used codon-based models of molecular evolution to examine how multiple variables related to ecology, habitat use, and light environment have influenced the evolution of flatfish RH1. We found signatures of positive selection across the flatfish phylogeny: a strong signal of positive selection in migratory species, and a pattern of increasingly positive selection in RH1 with increasing habitat depth. Furthermore, many flatfish lineages show evidence of RH1 spectral tuning mutations that indicate adaptation to the different light environments. Overall, our findings confirm the importance of visual habitat in shaping the diversity of flatfish RH1.

## 2. METHODS

### 2.1 Genetic and ecological data assembly

Rhodopsin gene coding sequences (CDS) were collected from 173 species of flatfishes, including representatives of all 16 pleuronectiform families (Table 1). We used the R package PhylotaR to retrieve all flatfish RH1 sequences published in GenBank, totalling 165 species (R core team 2023, Wang et al. 2013, Clark et al. 2016). We included seven additional flatfish species by manually identifying and extracting RH1 genes from unannotated genomes published by Lu et al. (2021), using local BLAST (BLAST+, Camacho et al. 2008). Finally, we used PCR to amplify and sequence full RH1 sequences for three additional Achiridae species (PCR conditions provided in Table S1). Full details of all species included and accession numbers are listed in Table S2.

**Table 1.**
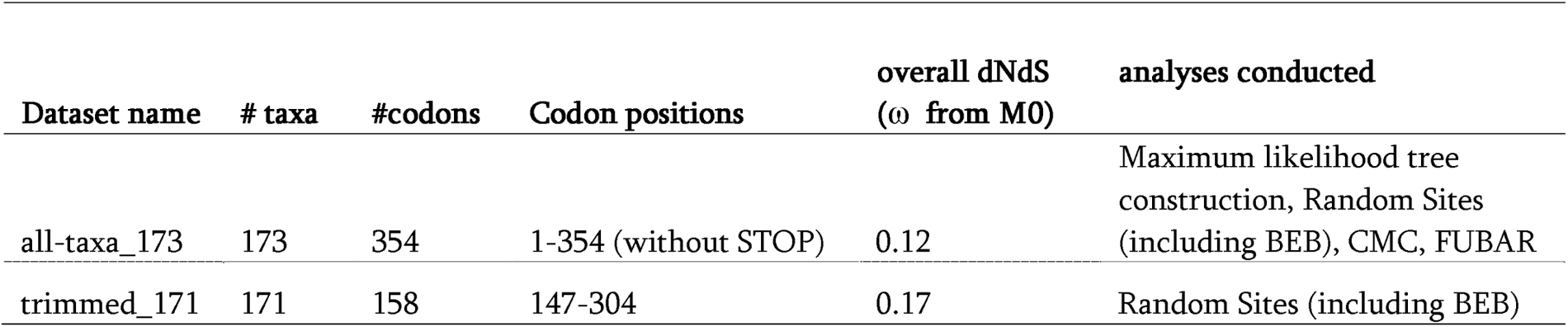
Dataset properties.

| Dataset name | # taxa | #codons | Codon positions | overall dNdS<br>($\omega$ from M0) | analyses conducted |
| --- | --- | --- | --- | --- | --- |
| all-taxa_173 | 173 | 354 | 1-354 (without STOP) | 0.12 | Maximum likelihood tree construction, Random Sites (including BEB), CMC, FUBAR |
| trimmed_171 | 171 | 158 | 147-304 | 0.17 | Random Sites (including BEB) |

A total of 173 RH1 sequences were aligned in AliView using MUSCLE codon alignment (Larsson 2014), and trimmed to remove stop codons. This alignment *“all-taxa_173”* included partial (∼474bp) and complete (1056bp) coding sequences collected for all species, and was used for most molecular evolutionary analyses (Table 1). We also generated a reduced alignment (*“trimmed_171”*) which consisted of only the 474 RH1 nucleotides available for 171 species; this alignment was used to validate results of the random-sites analyses and determination of positively selected sites obtained via the complete dataset *all-taxa_173* (see below for more details). We estimated a maximum likelihood gene tree using the *all-taxa_173* dataset with aLRT SH-like branch supports in PhyML 3.0 (Table 1, Figure S1; Guindon et al. 2010). This topology was used for all subsequent molecular evolutionary analyses (with both the *all-taxa_173* and *trimmed_171* datasets).

We collected data for four categories of ecological variables that we hypothesized would be most relevant for shaping flatfish rhodopsin evolution based on past research on fish visual systems. For example, adaptation of RH1 expression and structure in response to migrational movement (Hope et al. 1998, Wang et al. 2014) and evolutionary transitions to different habitat depths (Hunt et al. 1996, Dungan et al. 2015) have been described. Marine-freshwater evolutionary transitions have been demonstrated to affect functional evolution of RH1 (Van Nynatten et al. 2015, Van Nynatten et al. 2021), while different feeding modes have also been associated with adaptation in the visual system in flatfishes (Matsuda et a. 2008).

Data related to migratory behaviour, habitat depth, freshwater/marine classifications, and diet for flatfish species was gathered from FishBase using the R package rfishbase 4.0 (Froese and Pauly 2000, Boettinger et al. 2012). Migration was categorized as the presence/absence of oceanodromy. Species were classified into ocean zone-based depth categories: epipelagic (0-200m), mesopelagic (200-1000m), and bathypelagic (1000m+). Habitat was categorized as either marine or freshwater/brackish. DeGroot (1971) described three main feeding categories for flatfishes: (1) visual feeders of free-swimming prey; (2) visual feeders of benthic prey; (3) non-visual feeders of benthic prey. We combined (1) and (2) to form two categories: visual feeders and non-visual feeders. Diet information was obtained from FishBase, and we also searched the literature for additional diet data (Gregory 1933, de Groot 1969, Yazdani 1969, de Groot 1971, Holmes and Gibson 1983, Livingston 1987, Gibb 1997, Gibson 2008). Our categorizations for each species are provided in Table S2.

### 2.2 Analyses of molecular evolution in flatfish RH1

To investigate patterns of natural selection in RH1 across the flatfish phylogeny, we used codon-based molecular evolutionary models which estimate the ratio of non-synonymous-to-synonymous nucleotide substitution rates (dN/dS, or ω). When ω = 1, dN and dS are approximately equal and the gene is considered to be neutrally evolving. Values of ω <1 are indicative of purifying selection, while values of ω >1 are indicative of positive or diversifying selection (Yang 2007). Protein-coding genes are typically under purifying selection to preserve function, and elevated ω can therefore indicate adaptive evolution (Chang et al. 2012). If elevated dN/dS correlates with certain ecological attributes, it may suggest that adaptive evolution is occurring through ecologically-mediated selection pressures (Chang et al. 2012).

The random-sites model from the PAML packages’ codeml program was used to estimate dN/dS across the flatfish alignment (Yang 2007). Random-sites analyses involve fitting the alignment and phylogenetic tree to eight models of molecular evolution with differing parameters (see Table 2), and comparing the fits of certain pairs using *ln likelihood* values (lnL) in a likelihood ratio test (LRT). Significant chi-squared distribution statistics, considering the LRT and model complexity, are regarded as overall evidence of positive selection in the gene (Yang 2007). We applied random-sites analyses to our two datasets, *all-taxa_173* and *trimmed_171* to test for overall evidence of positive selection in flatfish RH1.

**Table 2.**
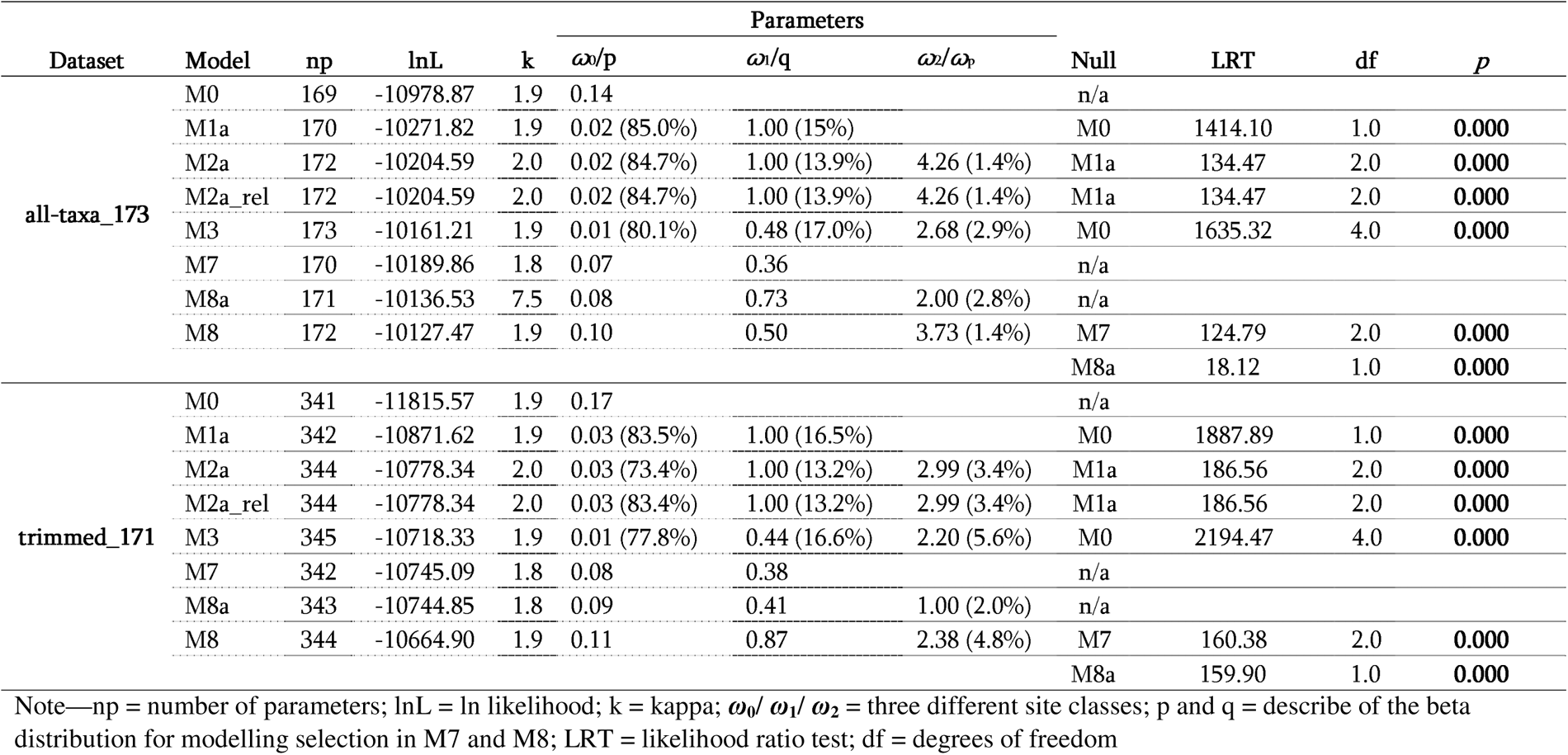
LRTs for Random-Sites Models in PAML for the flatfish RH1 gene tree.

We used PAML’s Clade Model C (CMC) to test whether four key aspects of flatfish ecology impacted the evolution of RH1. This model tests for differences in evolutionary rate between user-defined foreground “test” or “foreground” branches hypothesized to be experiencing differential selective pressures relative to background branches (Yang 2007). The significance of each model was calculated using LRTs compared to the null model *M2a_rel*. Non-nested CMC models were assessed for best fit using Akaike Information Criterion (AIC). Our CMC tests considered the effects of migration (oceanodromy vs. non-migration), depth (epipelagic vs. mesopelagic vs. bathypelagic), habitat (marine vs. freshwater/brackish) and diet (visual vs. non-visual feeding).

We used HyPhy’s RELAX model to test whether species with reduced reliance on visual cues (i.e., deepwater species and non-visual feeders) may have experienced a relaxation of selection on RH1. Similar to CMC, RELAX estimates ω among three rate classes, but fits an additional parameter estimating the strength of selection (*k*). ω values are transformed by *k* (ω*^k^*), wherein *k* > 1 pushes site classes with high or low ω values away from 1 (suggesting an intensification of selection); conversely, *k* < 1 shifts ω rate classes with high or low ω values towards 1 (suggesting relaxed selection; Wertheim et al. 2015). A model estimating a single *k* for all branches is compared against a model estimating *k* for the two branch classes corresponding to test (foreground) and reference (background) lineages in the relax analyses.

### 2.3 Site-by-site selection patterns

HyPhy’s FUBAR was used to compute a ω value for every codon in the gene alignment (Murrell et al. 2013; Table 3). Individual sites under positive selection were also inferred from the Bayes Empirical Bayes (BEB) output from the random sites analysis using the M8 model in PAML (Yang 2007, Table 3). The RH1 alignment and tree was also examined for evidence of amino acid changes at sites known to be associated with changes in RH1 function (e.g., spectral tuning and activation kinetics; Table 3; Hunt et al. 1996, Dungan and Chang 2017).

**Table 3.**
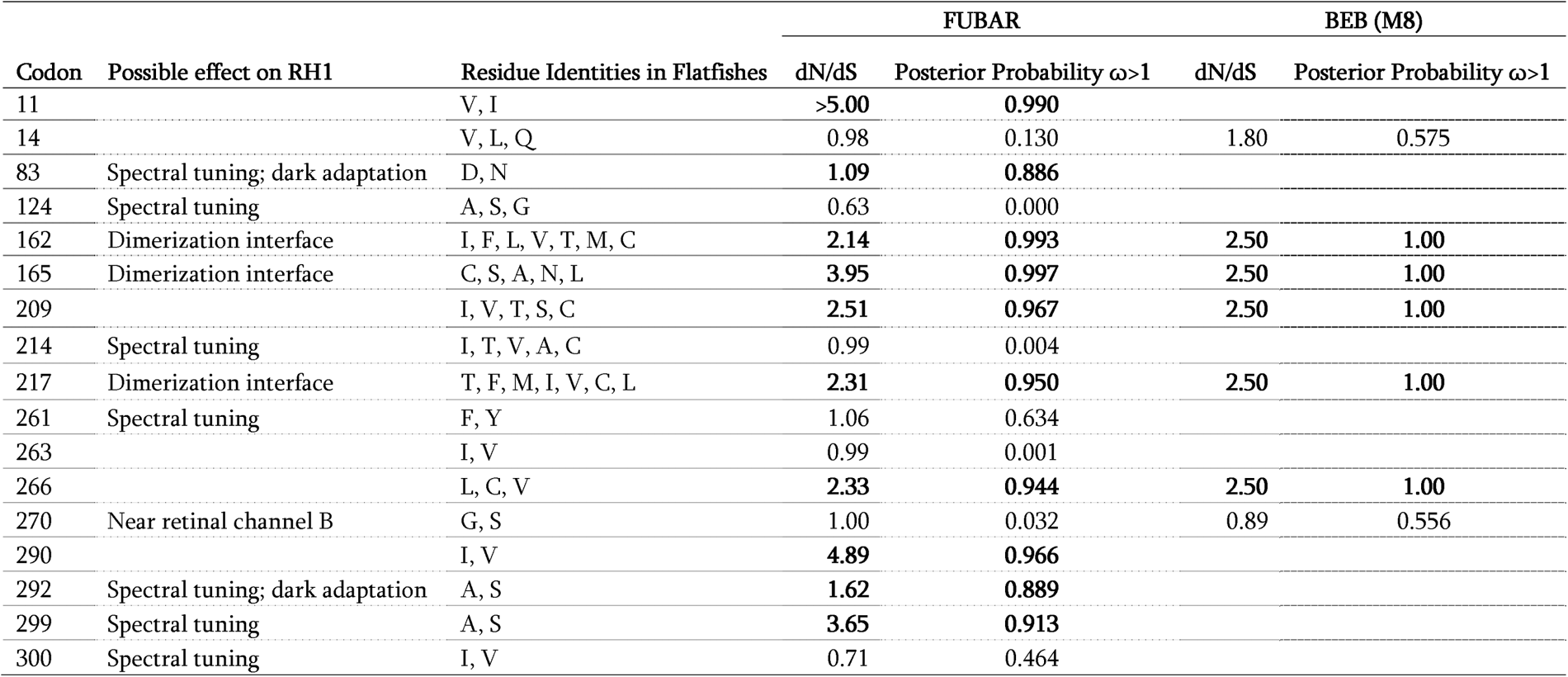
Rhodopsin residues discussed in text, including sites identified as under positive selection, and other sites of interest.

To visualize specific genetic changes in the opsin protein structure, we created two models of flatfish RH1 and mapped all sites of interest. We used the R package ggplot2 to generate a 2D snakeplot showing the opsin protein sequence and secondary structures (Wickam 2013). We used the homology model feature in ChimeraX ver. 1.16 from the MODELLER plug-in to model a representative species *Hippoglossus hippoglossus* (Atlantic halibut) RH1 protein sequence using the Bovine dimeric crystal RH1 structure (PDB1u19; Pettersen et al. 2004) as a reference. All proceeding codon numbering uses bovine RH1 sites.

## 3. Results

### 3.1 Phylogeny/random sites

We assessed the accuracy of our RH1 gene tree by comparing relationships to established flatfish phylogenies, and determined that the topology is largely congruent with previously published phylogenetic trees (Betancur-R et al. 2013, Campbell et al. 2019). Since several flatfish family relationships remain contentious, we decided that using the RH1 gene tree for molecular evolutionary analyses was the most conservative approach. This topology was used for all analyses.

Our analyses with PAML random sites found evidence for positive selection in flatfish RH1 in both the *all-taxa_173* dataset (ω = 3.73, M8 vs. M8a p <0.0001) and the *trimmed_171* alignment (ω = 2.38, M8 vs. M8a p < 0.0001) (Table 2), and as such further investigations of ecology-mediated and site-by-site selection patterns use the more complete *all-taxa_173* dataset.

### 3.2 Ecologically-mediated shifts in selection in flatfish RH1

We tested whether flatfish RH1 experienced differential selection pressures in response to a diverse array of ecological variables using CMC in PAML. Out of the 5 models, all 5 were significantly better fitting than the null M2a_rel, suggesting that all four aspects of life history and ecology tested (migration, depth, water type, and feeding behaviour) affect flatfish vision evolution (Table 4). The best-fitting model (identified with AIC) was one where migratory species were selected as “test” branches; these lineages showed strong evidence for positive selection (Figure 2, Table 4, ω_oceanodr._ = 10.48 / ω_BG_ = 2.89, p < .00). The multiple-partition depth model, isolating species in different ocean zones, identified an interesting pattern of increasing dN/dS with increasing habitat depth (ω_epi._ = 2.76 / ω_meso._ = 3.70 / ω_bathy._ = 6.22, p < .00, ΔAIC = 16.985). All partitions and AIC values are listed in Table 4, and are graphically represented in Figure 2.

**Figure 1.**
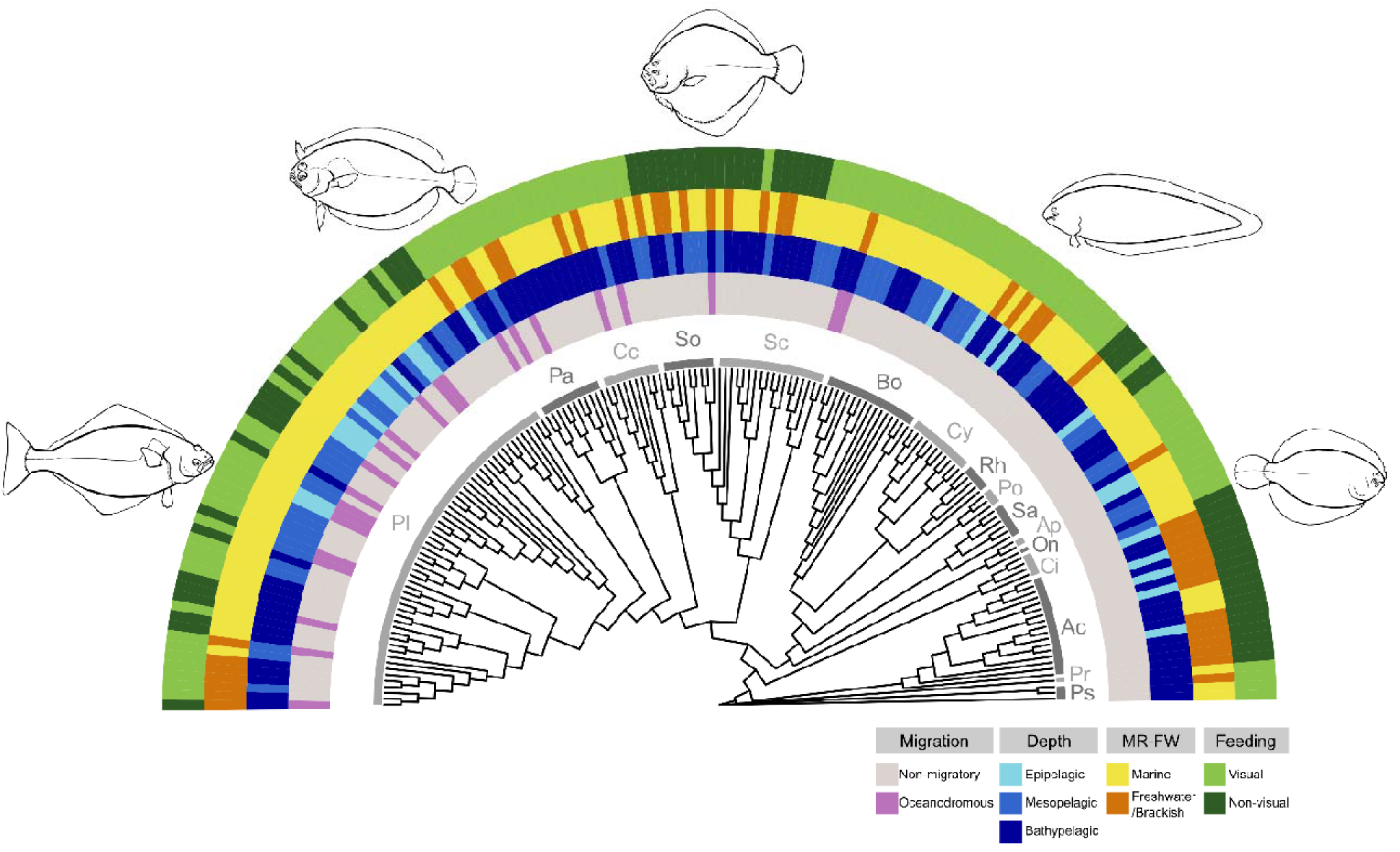
Flatfish RH1 gene tree topology, showing distribution of different ecological variables across the phylogeny that were used to inform PAML Clade Model C analysis partitions to investigate molecular evolution. Family name abbreviations: *Ps—*Psettodidae*, Pr—*Paralichthodidae*, Ac—*Achiridae*, Ci—* Citharidae*, On—*Oncopteridae*, Ap—*Achiropsettidae*, Sa—*Samaridae*, Po—*Poecilopsettidae*, Rh—* Rhombosoleidae*, Cy—*Cynoglossidae*, Bo—*Bothidae*, Sc—*Scophthalmidae*, So—*Soleidae*, Cc—* Cyclopsettidae*, Pa—*Paralichthyidae*, Pl—*Pleuronectidae.

**Figure 2.**
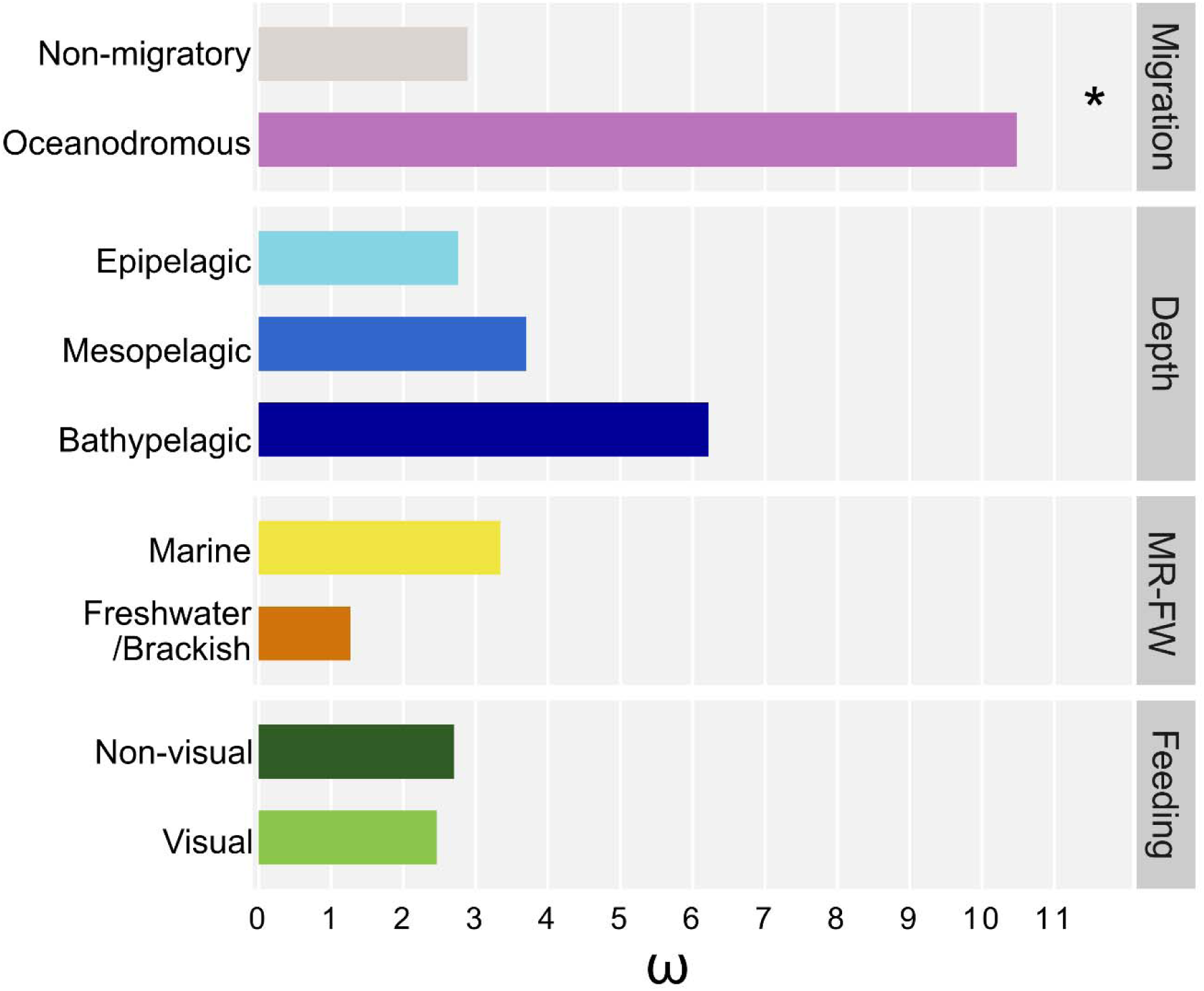
Bar plot showing ω values (divergent site class) of flatfish RH1 from four different clade model C analyses. Asterisk denotes the best-fitting model, as identified using Akaike Information Criterion.

**Table 4.**
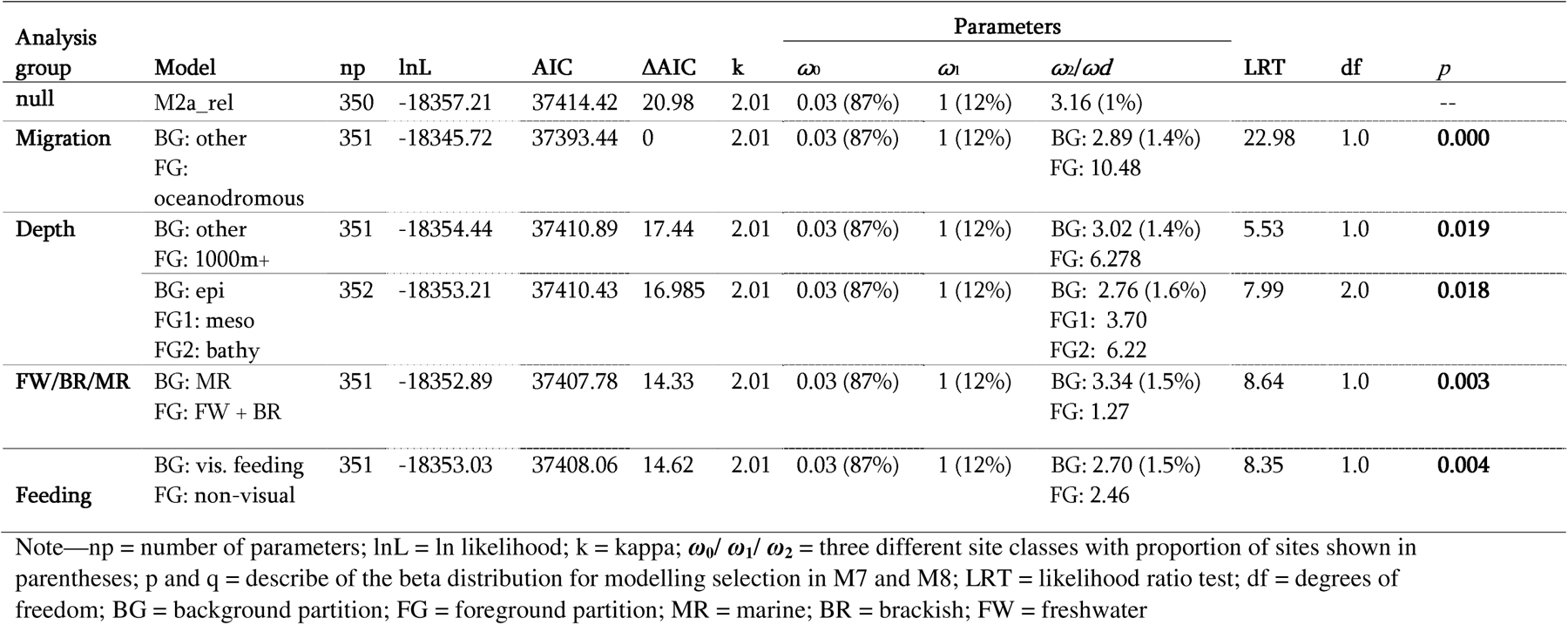
PAML Clade Model C test results for divergence between ecological partitions.

These results were further supported by analysis of the same four ecological categories using RELAX, which also determines shifts in the intensity of selection between selected “foreground” branches and the remaining “background” within a phylogeny. Overall, we identified significant intensification of selection in deepwater species (k = 1.34, P < .00, Table 5), consistent with the positive selection observed using CMC models. We also determined that non-visual feeders experienced significant relaxation (k = 0.86, p = .02), suggesting that the positive selection identified with CMC might be a result of decreases in the intensity of selection acting on RH1 in these lineages. Despite the strong positive selection observed in migratory and marine species, RELAX did not detect significant shifts in selection intensity for either of these partitions.

**Table 5:** Results from RELAX analyses for ecological partitions of flatfish RH1.

| Dataset | Test<br>(foreground)<br>branches | Reference<br>(background)<br>branches | Model | lnL | np | AIC | *k* | LR | *p* | Relaxation or<br>intensification<br>of selection |
| --- | --- | --- | --- | --- | --- | --- | --- | --- | --- | --- |
| all-taxa_173 | Oceanodromous | Non-migratory | Null | -19115.4 | 362 | 38959.1 | 1 | -- | -- | n.s. |
|  |  |  | Alternative | -19113.8 | 363 | 38958.0 | 1.19 | 3.13 | 0.077 |  |
|  | Bathypelagic<br>(1000m+) | Other | Null | -19162.0 | 362 | 39052.3 | 1 | -- | -- | intensification |
|  |  |  | Alternative | -19153.4 | 363 | 39037.1 | 1.34 | 17.21 | **0.000** |  |
|  | Non-visual<br>feeding | Visual<br>feeding | Null | -19115.3 | 362 | 38959.0 | 1 | -- | -- | relaxation |
|  |  |  | Alternative | -19112.6 | 363 | 38955.6 | 0.86 | 5.39 | **0.020** |  |
|  | Fresh/brackish | marine | Null | -19113.4 | 367 | 38965.2 | 1 | -- | -- | n.s. |
|  |  |  | Alternative | -19112.0 | 368 | 38964.5 | 0.85 | 2.70 | 0.100 |  |

Note—np = number of parameters; lnL = ln likelihood; AIC = Akaike information criterion; *k* = relaxation coefficient where *k*=1 is the null, *k* > 1 indicates intensification of selection, *k* < 1 indicates relaxation of selection; LR = likelihood ratio test of alternative and null models; df = degrees of freedom; *p* = p-value;

### 3.3 Variation at key functional sites in flatfish RH1

Analyses with FUBAR, which assess site-by-site dN/dS, revealed ten sites under pervasive positive selection and 249 sites under pervasive purifying selection, with a posterior probability cutoff of 0.8 (Figure 3, Table 3). Three of the positively selected sites (83, 292, 299) contains amino acid substitutions hypothesized to have a spectral tuning effects in fish RH1 (Dungan and Chang 2017, Hunt et al 1996, Yokoyama et al. 1995). The BEB analysis from the PAML M8 model indicated eight sites under significant positive selection, four of which were the same as in the FUBAR analysis. (Figure 3, Table 3). Several additional known spectral tuning sites (sites 124, 214, 261, and 300), while not under positive selection, exhibited possibly functionally relevant variation in flatfish. All amino acid identities found in flatfishes for sites of interest are shown in Table 3.

**Figure 3.**
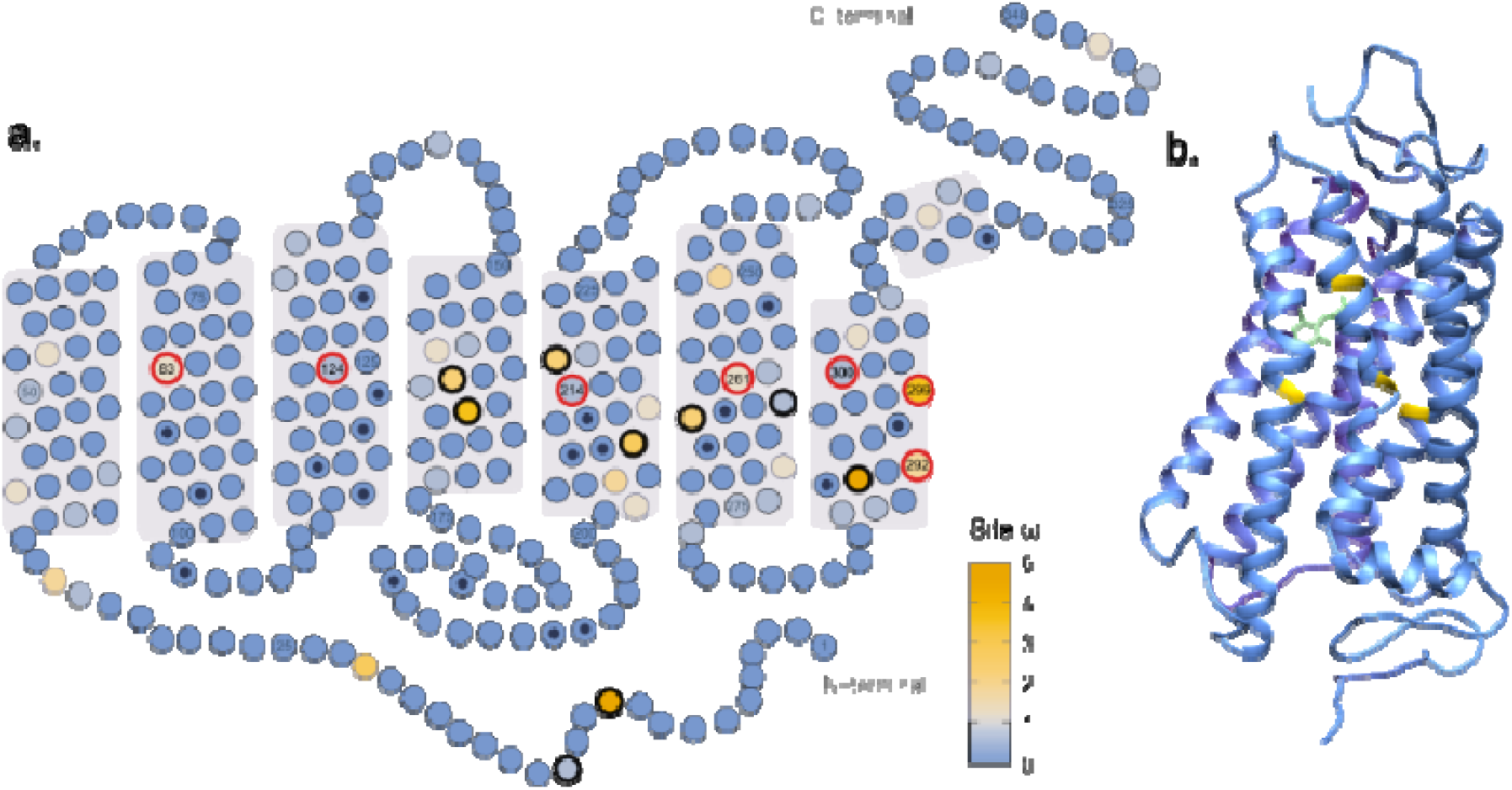
Flatfish RH1 snakeplot and protein homology model. ***A***—2D structure snakeplot showing the 7 transmembrane helices of the RH1 sequence, with sites coloured by FUBAR site ω value; highlighted residues with black borders are positively selected sites (combined FUBAR and BEB analyses); red borders denote variable spectral tuning sites in flatfishes. Black dots identify other known tuning sites in RH1 (from Musilova et al. 2021). ***B***—homology model of *Hippoglossus hippoglossus* (Atlantic halibut) sequence mapped to Bovine RH1 crystalized structure, showing proximity of retinal chromophore molecule (green) to the variable spectral tuning sites (yellow) identified in flatfishes.

## 4. DISCUSSION

### 4.1 Positive selection in RH1 across Flatfishes

Flatfishes are a highly species-rich order of benthic fish that inhabit a broad array of environments, ranging from deepwater marine communities to shallow coastal reefs to freshwater systems (Gibson 2008). This variability in aquatic habitats means flatfish have evolved in a rich diversity of lighting environments. We found evidence for positive selection in flatfish RH1, and that selection on this gene was strongly associated with ecology. Models associated with depth and migration fit the data better than diet-based tests, suggesting that molecular changes in RH1 are being driven more strongly by habitat light environment than by vision related to feeding. Specifically, RH1 evolved under positive selection in oceanodromous and deepwater flatfishes, and under purifying selection in freshwater species. In conjunction with these findings, we also identified several mutations across the flatfish phylogeny that likely result in functional adaptations in RH1. Overall, our findings demonstrate that the remarkable variation in flatfish RH1 is likely a result of ecologically-driven adaptation.

### 4.3 Strong positive selection in RH1 in oceanodromous flatfish

We found that oceanodromous flatfishes experienced significantly stronger positive selection in RH1 relative to their non-oceanodromous counterparts. Oceanodromous flatfishes partake in cyclical migration in a manner similar to tuna, wherein they undergo yearly movements through open ocean between feeding and spawning grounds (Hart, 1991, Gibson 2008). We used the description of oceanodromy provided by FishBase: substantial regular movements through open-ocean waters. There appears to be no correlation between oceanodromy and other ecological characters analyzed (see Figure 1); thus, oceanodromy appears to represent a distinct driver of RH1 evolution in these fishes.

The effect of oceanodromy on vision evolution may be reflective of adaptation driven by a need for functional vision under more diverse light environments than flatfish typically occupy (Guo et al. 2022). Flatfish are benthic specialists, with a high modified body plan suited for life on the ocean floor, and therefore experience a relatively consistent light environment (Zhang et al. 2022). Oceanodromous species that enter the open ocean would be exposed to brighter ambient light, potentially more or less blue-shifted depending on the depth they otherwise occupy (Hauser and Chang 2017). These migrations would likely impose selective pressure on RH1 to function optimally in both dim benthic and brighter open-ocean conditions, as opposed to the more consistent photic environments inhabited by their nonmigratory relatives. This signal of positive selection opens a promising avenue for further investigation. Future work could examine how other visual transduction genes have evolved in concert with RH1 to optimize the flatfish visual system to function in both dim and bright environments.

### 4.4 RH1 molecular evolutionary rates increase with habitat and depth

The relationship between depth and RH1 molecular evolution is well established, with deep-dwelling species often accumulating blue-shifting mutations to better match the ambient spectral content (Hunt et al. 1996, Dungan et al. 2016, Luk et al. 2021, Van Nynatten et al. 2021). Flatfishes occupy depths from <1 m to >3000 m below sea level, spanning the photic zone to well below the limit of light penetration underwater. Beyond a depth of 1000m, the only light visible is generated by bioluminescent organisms (Xia et al. 2021). We hypothesized that this extensive variation in habitat depth would impose differential selection pressures on flatfish RH1.

With two independent molecular evolutionary models, we identified a pattern of increased positive selection coinciding with increasing habitat depth, as well as increasing intensity of selection in deepwater species. These findings are similar to that observed by Dungan et al. (2016), who identified a pattern of positive selection in epipelagic and bathypelagic cetaceans relative to a lower ω in mesopelagic species, reflective of divergence away from a mesopelagic cetacean common ancestor. Based on fossil evidence, the flatfish common ancestor was likely similar to the sub-order Psettoidea, which contains three epipelagic species that comprise the sister group to all other flatfishes (Campbell et al. 2019). Furthermore, all proposed sister taxa to all the flatfishes are largely epipelagic or brackish-dwelling (including lates perches (Latidae), threadfins (Polynemidae), archerfish (Toxotidae), jacks (Carangidae), and Snooks (Centropomidae) suggesting that the flatfish common ancestor was also a shallow-water species (Bentacur et al. 2013, Campbell et al. 2019, Atta 2022). Flatfish RH1 may therefore exhibit a similar pattern of selection to what has been found in whales: RH1 diversification along the depth gradient may be associated with adaptation to progressively deeper environments. Ancestral flatfish RH1 would likely be tuned to function most effectively in the bright light of epipelagic zones. We hypothesize that adaptation to deeper waters was accompanied by strong positive selection on RH1 to function under increasingly darker and more blue wavelength-dominated environments. Future research of depth-based adaptation in RH1 could include comparative studies of RH1 function and expression between deep- and shallow-water flatfish species to illuminate functional differences and identify convergence among repeated deep-water transitions.

We found evidence that RH1 was under purifying selection in freshwater and brackish flatfishes. This result was unexpected given that transitions from marine to freshwater often result in positive selection in RH1 in other fish lineages, including anchovies and croakers (Van Nynatten et al. 2015, Van Nynatten et al. 2021). We found South American achirids were under particularly strong purifying selection. This result may be due to the small number of exclusively freshwater species included (n=6), the incomplete sequence data available for these species, or the fact that achirid species are recently diverged (Bitencourt et al. 2023) and therefore may not have accumulated substantial variation in RH1.

### 4.4 Spectral tuning mutations in flatfish RH1

To analyze selection patterns across amino acid sites in the flatfish RH1 gene, we first used two computational models (BEB & FUBAR) to detect positively selected sites and examined specific amino acid residues at these sites. Of particular interest were spectral tuning sites, where known amino acid changes are hypothesized to alter the λ_max_ of visual opsins and contribute to adaptation to ambient light conditions (Yokoyama et al. 1995, Lin et al. 2017, Hauser and Chang 2017). We identified several interesting correlations between residue distributions and ecological characteristics in flatfish RH1 genes, outlined below.

The D83N substitution, found in many flatfishes, mediates a small blue-shift in teleosts, which is considered adaptive for fishes inhabiting deep water, or other such dim environments with a greater prevalence of blue-wavelength light (Hauser et al. 2017). Interestingly, the N83 residue is predominant in flatfishes, and occurs in 106 of the 137 species for which site has been sequenced; D83 is found only in migratory, freshwater, or shallow-water flatfish species. Hauser et al. (2017) performed site-directed mutagenesis and determined that in addition to shifting the protein’s λ_max_, site 83 variation can affect kinetics and RH1 recovery post-activation. Because RH1 with site N83 has a significantly longer recovery and greater sensitivity, it is likely beneficial in dim light environments; we propose that the widespread N83 residue in flatfishes is adaptive for their dim and invariable benthic habitats, especially in non-migratory and deepwater species.

Transitions between A and S residues at two sites in RH1 also cause shifts in λ_max_in teleosts. At both sites 292 and 299, A is slightly more common in teleosts, with mutations to S causing a large blue-shift in S292 and small red-shift in S299 (Hunt et al. 1996). Mutations between A and S at 292 and 299 in RH1 have been frequently associated with habitat transitions in fishes, especially changes in depth. In flatfishes we identified A292/S299 almost exclusively in freshwater and epipelagic species, aligning with expectations of red-shifted RH1 in species dwelling in red-shifted freshwater and bright shallower habitats. Additionally, the deepest-dwelling meso- and bathypelagic species show either S292 or A299 (one of the two blue-shifting residues), again consistent with adaptations to blue-shifted deep marine environments.

In flatfishes, RH1 site 261 exhibits a pattern of depth-based transitions between F261 (common in teleosts) and the red-shifting Y261 (Yokoyama et al. 1995, Hunt et al 1996, Hill et al. 2019). All flatfishes with Y261 were epipelagic or shallow mesopelagic (maximum depth∼300m), consistent with previous findings associating the F261Y mutation with shallow-dwelling fishes. Interestingly the only freshwater species from the marine family Cynoglossidae (tonguefish) has both A292/S299 and has independently evolved Y261, suggesting independent adaptation to the longer wavelengths typically found in freshwater.

### 4.5 Conclusions

We found strong support for ecologically-mediated diversification in flatfish RH1, as well as parallel amino acid substitutions in RH1 to promote functional convergence to different environmental conditions. This means many indicators of adaptation to deepwater are independent of each other (Campbell et al. 2019). We identified trends in RH1 evolutionary rate that connect the ancestral shallow-water marine flatfishes to more recent deepwater adaptations, showing that deep-dwelling flatfishes have undergone more rapid evolution in response to the changes in aquatic light conditions. Selection at sites associated with RH1 spectral tuning also highlight the history of visual system modifications shared by flatfishes undergoing habitat transitions, and show repeated convergence towards a more blue- or red-shifted λ_max_ depending on current environmental conditions. RH1 in flatfishes is a highly variable gene and has experienced selection in response to several key changes in flatfish visual ecology. The diversity of flatfishes and their unique specializations to a variety of aquatic habitats make them a promising model for future investigations of visual ecology and molecular adaptation to different environments.

## DATA AVAILABILITY

The rhodopsin sequence data that support the findings of this study are available in GenBank at https://www.ncbi.nlm.nih.gov/genbank/ (and see Table S2).

## ACKNOWLEDGEMENTS

UTEA to EM, NSERC PDF and UTSC PDF to FEH, MITACS to AVN, BSWC Discovery Grant, NRL Discovery Grant. We thank Anthony Rajkumar for comments on an earlier version of this manuscript, and Lisa Byrne for valuable discussions.

