## Supplemental Tables and figures for "Evolution of rhodopsin in flatfishes (Pleuronectiformes) is associated with depth and migratory behaviour"

**Table S1.** PCR program and primers used to amplify and sequence RH1 for this study

|  | Step | | Temp | Time |
| --- | --- | --- | --- | --- |
| Thermocycler Protocol | 1) Initial melting | | 94°C | 4 min |
|  | 2) Melting | | 95°C | 30 sec |
|  | 3) Annealing | | 52°C | 30 sec |
|  | 4) Elongation | | 72°C | 30 sec |
|  | *Repeat steps 2-4 35x* | | | |
|  | 5) Final elongation | | 72°C | 5 min |
| F primer | Rh1 193F | From Chen et al. 2003 | | |
| R primer | Rh1 1039R | From Chen et al. 2003 | | |

**Table S2.** Species information for RH1 sequences, and ecological data used for Clade Model C (CMC) partitions. * indicates sequences generated for this study.

| Family | Species | GenBank  Accession # | Migration | Habitat Depth | Water type  (Fresh/Brackish/  Marine) | Feeding Type |
| --- | --- | --- | --- | --- | --- | --- |
| Achiridae | *Gymnachirus melas* | JQ938030.1 | Non-migratory | Bathypelagic | MR | Non-visual |
| Achiridae | *Gymnachirus nudus* | ON562520.1 | Non-migratory | Bathypelagic | MR | Non-visual |
| Achiridae | *Gymnachirus texae* | JQ938031.1 | Non-migratory | Bathypelagic | MR | Non-visual |
| Achiridae | *Apionichthys finis* | ON562486.1 | Non-migratory | Freshwater | FW | Non-visual |
| Achiridae | *Catathyridium jenynsii* | ON562493.1 | Non-migratory | Freshwater | FW | Non-visual |
| Achiridae | *Hypoclinemus mentalis* | AV201720 * | Non-migratory | Freshwater | FW | Non-visual |
| Achiridae | *Hypoclinemus sp.* | JQ938065.1 | Non-migratory | Freshwater | FW | Non-visual |
| Achiridae | *Achirus declivis* | ON562472.1 | Non-migratory | Bathypelagic | BR | Non-visual |
| Achiridae | *Achirus lineatus* | AV201701 * | Non-migratory | Bathypelagic | BR | Non-visual |
| Achiridae | *Achirus mucuri* | ON562483.1 | Non-migratory | Epipelagic | BR | Non-visual |
| Achiridae | *Trinectes inscriptus* | ON562503.1 | Non-migratory | Bathypelagic | BR | Non-visual |
| Achiridae | *Trinectes microphthalmus* | ON562507.1 | Non-migratory | Bathypelagic | BR | Non-visual |
| Achiridae | *Achirus achirus* | AV201702 * | Non-migratory | Bathypelagic | FW/BR | Non-visual |
| Achiridae | *Apionichthys dumerili* | ON562489.1 | Non-migratory | Bathypelagic | FW/BR | Non-visual |
| Achiridae | *Catathyridium garmani* | ON562495.1 | Non-migratory | Bathypelagic | FW/BR | Non-visual |
| Achiridae | *Trinectes maculatus* | JAFBJQ010001709.1 | Non-migratory | Bathypelagic | FW/BR | Non-visual |
| Achiridae | *Trinectes paulistanus* | ON562515.1 | Non-migratory | Bathypelagic | FW/BR | Non-visual |
| Achiropsettidae | *Mancopsetta maculata* | KF312129.1 | Non-migratory | Epipelagic | MR | Visual |
| Achiropsettidae | *Neoachiropsetta milfordi* | JQ938075.1 | Non-migratory | Epipelagic | MR | Visual |
| Bothidae | *Arnoglossus imperialis* | JQ938032.1 | Non-migratory | Mesopelagic | MR | Visual |
| Bothidae | *Arnoglossus laterna* | EU224095.1 | Non-migratory | Mesopelagic | MR | Visual |
| Bothidae | *Asterorhombus cocosensis* | JQ938033.1 | Non-migratory | Bathypelagic | MR | Visual |
| Bothidae | *Bothus ocellatus* | ON562522.1 | Non-migratory | Bathypelagic | MR | Visual |
| Bothidae | *Bothus podas* | AY368313.1 | Non-migratory | Mesopelagic | MR | Visual |
| Bothidae | *Bothus robinsi* | JQ938034.1 | Non-migratory | Bathypelagic | MR | Visual |
| Bothidae | *Chascanopsetta lugubris* | JAFBJT010105763.1 | Non-migratory | Epipelagic | MR | Visual |
| Bothidae | *Japonolaeops dentatus* | MK617114.1 | Non-migratory | Mesopelagic | MR | Visual |
| Bothidae | *Kamoharaia megastoma* | MK617115.1 | Non-migratory | Mesopelagic | MR | Visual |
| Bothidae | *Laeops kitaharae* | JQ938035.1 | Non-migratory | Mesopelagic | MR | Visual |
| Bothidae | *Monolene sp.* | JQ938072.1 | Non-migratory | Mesopelagic | MR | Visual |
| Bothidae | *Neolaeops microphthalmus* | MK617117.1 | Non-migratory | Mesopelagic | MR | Visual |
| Bothidae | *Parabothus taiwanensis* | MK617118.1 | Non-migratory | Epipelagic | MR | Visual |
| Bothidae | *Psettina tosana* | JQ938036.1 | Non-migratory | Mesopelagic | MR | Visual |
| Bothidae | *Taeniopsetta ocellata* | MK617126.1 | Non-migratory | Mesopelagic | MR | Visual |
| Bothidae | *Trichopsetta ventralis* | JQ938037.1 | Non-migratory | Bathypelagic | MR | Visual |
| Citharidae | *Brachypleura novaezeelandiae* | KF312132.1 | Non-migratory | Bathypelagic | MR | Visual |
| Citharidae | *Citharoides macrolepis* | KF312133.1 | Non-migratory | Mesopelagic | MR | Visual |
| Citharidae | *Citharus linguatula* | EF439360.1 | Non-migratory | Mesopelagic | MR | Visual |
| Citharidae | *Lepidoblepharon ophthalmolepis* | KF312135.1 | Non-migratory | Mesopelagic | MR | Non-visual |
| Cyclopsettidae | *Citharichthys arctifrons* | JQ938042.1 | Non-migratory | Mesopelagic | MR | Visual |
| Cyclopsettidae | *Citharichthys arenaceus* | ON562523.1 | Non-migratory | Bathypelagic | MR | Visual |
| Cyclopsettidae | *Citharichthys minutus* | MK617108.1 | Non-migratory | Bathypelagic | MR | Visual |
| Cyclopsettidae | *Citharichthys sp.* | MK617109.1 | Non-migratory | Bathypelagic | MR | Visual |
| Cyclopsettidae | *Cyclopsetta chittendeni* | JQ938043.1 | Non-migratory | Bathypelagic | MR | Visual |
| Cyclopsettidae | *Etropus microstomus* | JQ938045.1 | Oceanodromous | Bathypelagic | MR | Visual |
| Cyclopsettidae | *Syacium micrurum* | AY368334.1 | Non-migratory | Mesopelagic | MR | Visual |
| Cyclopsettidae | *Cyclopsetta querna* | MK617110.1 | Non-migratory | Bathypelagic | BR | Visual |
| Cyclopsettidae | *Etropus crossotus* | JQ938044.1 | Oceanodromous | Bathypelagic | BR | Visual |
| Cyclopsettidae | *Citharichthys gilberti* | MK617107.1 | Non-migratory | Bathypelagic | FW/BR | Visual |
| Cynoglossidae | *Cynoglossus browni* | DQ197839.1 | Non-migratory | Bathypelagic | MR | Visual |
| Cynoglossidae | *Cynoglossus nigropinnatus* | MK617112.1 | Non-migratory | Bathypelagic | MR | Visual |
| Cynoglossidae | *Paraplagusia japonica* | KF312136.1 | Non-migratory | Bathypelagic | MR | Visual |
| Cynoglossidae | *Symphurus atricauda* | JQ938040.1 | Non-migratory | Mesopelagic | MR | Visual |
| Cynoglossidae | *Symphurus hondoensis* | MK617124.1 | Non-migratory | Mesopelagic | MR | Visual |
| Cynoglossidae | *Symphurus orientalis* | KF312137.1 | Non-migratory | Bathypelagic | MR | Visual |
| Cynoglossidae | *Symphurus plagusia* | JQ938066.1 | Non-migratory | Bathypelagic | MR | Visual |
| Cynoglossidae | *Cynoglossus feldmanni* | MK617111.1 | Non-migratory | Freshwater | FW | Visual |
| Cynoglossidae | *Paraplagusia blochii* | JAFBJS010055197.1 | Non-migratory | Bathypelagic | BR | Visual |
| Cynoglossidae | *Cynoglossus semilaevis* | CM002382.1 | Non-migratory | Freshwater | FW | Visual |
| Cynoglossidae | *Cynoglossus senegalensis* | DQ197840.1 | Non-migratory | Bathypelagic | FW/BR | Visual |
| Oncopteridae | *Oncopterus darwinii* | JQ938081.1 | Non-migratory | Bathypelagic | MR | Visual |
| Paralichthodidae | *Paralichthodes algoensis* | MK617119.1 | Non-migratory | Bathypelagic | MR | Visual |
| Paralichthyidae | *Ancylopsetta ommata* | JQ938041.1 | Non-migratory | Bathypelagic | MR | Visual |
| Paralichthyidae | *Paralichthys albigutta* | JQ938071.1 | Non-migratory | Bathypelagic | MR | Visual |
| Paralichthyidae | *Paralichthys olivaceus* | HQ413772.1 | Oceanodromous | Mesopelagic | MR | Visual |
| Paralichthyidae | *Pseudorhombus dupliciocellatus* | JAFBJW010000168.1 | Non-migratory | Bathypelagic | MR | Visual |
| Paralichthyidae | *Pseudorhombus oligodon* | KF312138.1 | Non-migratory | Bathypelagic | MR | Visual |
| Paralichthyidae | *Pseudorhombus pentophthalmus* | JQ938046.1 | Non-migratory | Bathypelagic | MR | Visual |
| Paralichthyidae | *Xystreurys liolepis* | KF312139.1 | Non-migratory | Bathypelagic | MR | Visual |
| Paralichthyidae | *Paralichthys aestuarius* | MH032338.1 | Non-migratory | Bathypelagic | BR | Visual |
| Paralichthyidae | *Paralichthys californicus* | MH032339.1 | Oceanodromous | Bathypelagic | BR | Visual |
| Paralichthyidae | *Paralichthys dentatus* | KU980166.1 | Oceanodromous | Bathypelagic | BR | Visual |
| Paralichthyidae | *Paralichthys woolmani* | MH032342.1 | Non-migratory | Bathypelagic | BR | Visual |
| Pleuronectidae | *Acanthopsetta nadeshnyi* | MH032247.1 | Non-migratory | Mesopelagic | MR | Visual |
| Pleuronectidae | *Atheresthes evermanni* | MH032249.1 | Non-migratory | Epipelagic | MR | Visual |
| Pleuronectidae | *Atheresthes stomias* | MH032251.1 | Non-migratory | Mesopelagic | MR | Visual |
| Pleuronectidae | *Cleisthenes pinetorum* | MH032252.1 | Non-migratory | Mesopelagic | MR | Visual |
| Pleuronectidae | *Clidoderma asperrimum* | MH032256.1 | Non-migratory | Epipelagic | MR | Visual |
| Pleuronectidae | *Dexistes rikuzenius* | MH032259.1 | Non-migratory | Mesopelagic | MR | Visual |
| Pleuronectidae | *Embassichthys bathybius* | JQ938048.1 | Non-migratory | Epipelagic | MR | Visual |
| Pleuronectidae | *Eopsetta grigorjewi* | MH032263.1 | Non-migratory | Epipelagic | MR | Visual |
| Pleuronectidae | *Eopsetta jordani* | MH032265.1 | Non-migratory | Mesopelagic | MR | Visual |
| Pleuronectidae | *Glyptocephalus cynoglossus* | EU492060.1 | Oceanodromous | Epipelagic | MR | Non-visual |
| Pleuronectidae | *Glyptocephalus kitaharae* | MH032272.1 | Non-migratory | Mesopelagic | MR | Visual |
| Pleuronectidae | *Glyptocephalus stelleri* | MH032275.1 | Oceanodromous | Epipelagic | MR | Non-visual |
| Pleuronectidae | *Glyptocephalus zachirus* | MH032277.1 | Non-migratory | Mesopelagic | MR | Visual |
| Pleuronectidae | *Hippoglossoides dubius* | MH032280.1 | Oceanodromous | Mesopelagic | MR | Visual |
| Pleuronectidae | *Hippoglossoides elassodon* | MH032281.1 | Oceanodromous | Epipelagic | MR | Visual |
| Pleuronectidae | *Hippoglossoides platessoides* | MH032287.1 | Oceanodromous | Epipelagic | MR | Visual |
| Pleuronectidae | *Hippoglossus hippoglossus* | AF156265.1 | Oceanodromous | Epipelagic | MR | Visual |
| Pleuronectidae | *Hippoglossus stenolepis* | MH032291.1 | Oceanodromous | Epipelagic | MR | Visual |
| Pleuronectidae | *Isopsetta isolepis* | MH032292.1 | Oceanodromous | Mesopelagic | MR | Visual |
| Pleuronectidae | *Lepidopsetta bilineata* | MH032294.1 | Non-migratory | Mesopelagic | MR | Visual |
| Pleuronectidae | *Lepidopsetta mochigarei* | MH032296.1 | Non-migratory | Bathypelagic | MR | Non-visual |
| Pleuronectidae | *Lepidopsetta polyxystra* | MH032299.1 | Non-migratory | Mesopelagic | MR | Visual |
| Pleuronectidae | *Limanda aspera* | MH032302.1 | Non-migratory | Mesopelagic | MR | Non-visual |
| Pleuronectidae | *Limanda limanda* | MH032306.1 | Oceanodromous | Bathypelagic | MR | Visual |
| Pleuronectidae | *Limanda sakhalinensis* | MH032307.1 | Non-migratory | Mesopelagic | MR | Visual |
| Pleuronectidae | *Lyopsetta exilis* | JQ938073.1 | Non-migratory | Mesopelagic | MR | Visual |
| Pleuronectidae | *Microstomus achne* | MH032320.1 | Non-migratory | Mesopelagic | MR | Non-visual |
| Pleuronectidae | *Microstomus kitt* | MH032326.1 | Oceanodromous | Mesopelagic | MR | Non-visual |
| Pleuronectidae | *Microstomus pacificus* | MH032327.1 | Non-migratory | Epipelagic | MR | Non-visual |
| Pleuronectidae | *Microstomus shuntovi* | MH032329.1 | Non-migratory | Mesopelagic | MR | Non-visual |
| Pleuronectidae | *Myzopsetta ferruginea* | MH032332.1 | Non-migratory | Bathypelagic | MR | Visual |
| Pleuronectidae | *Myzopsetta proboscidea* | MH032333.1 | Non-migratory | Bathypelagic | MR | Visual |
| Pleuronectidae | *Myzopsetta punctatissima* | MH032337.1 | Non-migratory | Bathypelagic | MR | Non-visual |
| Pleuronectidae | *Nematops sp.* | MK617116.1 | Non-migratory | Mesopelagic | MR | Visual |
| Pleuronectidae | *Parophrys vetulus* | MH032343.1 | Oceanodromous | Mesopelagic | MR | Non-visual |
| Pleuronectidae | *Pleuronectes quadrituberculatus* | MH032356.1 | Non-migratory | Mesopelagic | MR | Visual |
| Pleuronectidae | *Pleuronichthys coenosus* | MH032357.1 | Non-migratory | Mesopelagic | MR | Non-visual |
| Pleuronectidae | *Pleuronichthys cornutus* | MH032359.1 | Non-migratory | Bathypelagic | MR | Non-visual |
| Pleuronectidae | *Pleuronichthys decurrens* | MH032362.1 | Non-migratory | Mesopelagic | MR | Non-visual |
| Pleuronectidae | *Pleuronichthys ritteri* | MH032369.1 | Non-migratory | Bathypelagic | MR | Visual |
| Pleuronectidae | *Pleuronichthys verticalis* | MH032371.1 | Non-migratory | Mesopelagic | MR | Non-visual |
| Pleuronectidae | *Psettichthys melanostictus* | MH032373.1 | Non-migratory | Mesopelagic | MR | Visual |
| Pleuronectidae | *Pseudopleuronectes americanus* | MH032374.1 | Oceanodromous | Bathypelagic | MR | Non-visual |
| Pleuronectidae | *Pseudopleuronectes herzensteini* | MH032377.1 | Non-migratory | Bathypelagic | MR | Non-visual |
| Pleuronectidae | *Pseudopleuronectes obscurus* | MH032381.1 | Non-migratory | Bathypelagic | MR | Non-visual |
| Pleuronectidae | *Pseudopleuronectes schrenki* | MH032383.1 | Non-migratory | Bathypelagic | MR | Visual |
| Pleuronectidae | *Pseudopleuronectes yokohamae* | LC586805.1 | Non-migratory | Bathypelagic | MR | Non-visual |
| Pleuronectidae | *Reinhardtius hippoglossoides* | MH032387.1 | Oceanodromous | Epipelagic | MR | Visual |
| Pleuronectidae | *Verasper moseri* | AB930176.1 | Non-migratory | Mesopelagic | MR | Visual |
| Pleuronectidae | *Verasper variegatus* | MH032392.1 | Non-migratory | Bathypelagic | MR | Non-visual |
| Pleuronectidae | *Hypsopsetta guttulata* | MH032365.1 | Non-migratory | Bathypelagic | BR | Visual |
| Pleuronectidae | *Pleuronectes platessa* | MH032354.1 | Oceanodromous | Mesopelagic | BR | Visual |
| Pleuronectidae | *Pleuronectes putnami* | MH032315.1 | Non-migratory | Bathypelagic | BR | Visual |
| Pleuronectidae | *Liopsetta glacialis* | MH032309.1 | Non-migratory | Bathypelagic | FW/BR | Visual |
| Pleuronectidae | *Liopsetta pinnifasciata* | MH032312.1 | Non-migratory | Bathypelagic | FW/BR | Visual |
| Pleuronectidae | *Platichthys bicoloratus* | MH032345.1 | Oceanodromous | Bathypelagic | FW/BR | Non-visual |
| Pleuronectidae | *Platichthys flesus* | MH032348.1 | Non-migratory | Bathypelagic | FW/BR | Visual |
| Pleuronectidae | *Platichthys stellatus* | MH032350.1 | Non-migratory | Mesopelagic | FW/BR | Visual |
| Poecilopsettidae | *Poecilopsetta beanii* | JQ938054.1 | Non-migratory | Epipelagic | MR | Non-visual |
| Poecilopsettidae | *Poecilopsetta plinthus* | KF312144.1 | Non-migratory | Mesopelagic | MR | Non-visual |
| Psettodidae | *Psettodes erumei* | JAFBJZ010000915.1 | Non-migratory | Bathypelagic | MR | Visual |
| Psettodidae | *Psettodes sp.* | AF148143.1 | Non-migratory | Bathypelagic | MR | Visual |
| Psettodidae | *Psettodes belcheri* | AY368332.1 | Non-migratory | Bathypelagic | BR | Visual |
| Rhombosoleidae | *Colistium nudipinnis* | JAFBJU010000782.1 | Non-migratory | Bathypelagic | MR | Visual |
| Rhombosoleidae | *Peltorhamphus latus* | JX049180.1 | Non-migratory | Bathypelagic | MR | Non-visual |
| Rhombosoleidae | *Rhombosolea leporina* | MK617123.1 | Non-migratory | Bathypelagic | MR | Non-visual |
| Rhombosoleidae | *Rhombosolea plebeia* | JQ938068.1 | Non-migratory | Bathypelagic | MR | Visual |
| Rhombosoleidae | *Rhombosolea tapirina* | JQ938069.1 | Non-migratory | Bathypelagic | BR | Non-visual |
| Samaridae | *Plagiopsetta glossa* | JQ938056.1 | Non-migratory | Mesopelagic | MR | Visual |
| Samaridae | *Plagiopsetta sp.* | MK617121.1 | Non-migratory | Mesopelagic | MR | Visual |
| Samaridae | *Samariscus japonicus* | JQ938057.1 | Non-migratory | Bathypelagic | MR | Visual |
| Samaridae | *Samariscus latus* | KF312146.1 | Non-migratory | Bathypelagic | MR | Visual |
| Samaridae | *Samariscus xenicus* | JQ938058.1 | Non-migratory | Bathypelagic | MR | Visual |
| Samaridae | *Samaris cristatus* | KF312145.1 | Non-migratory | Bathypelagic | BR | Visual |
| Scophthalmidae | *Lepidorhombus boscii* | EF439123.1 | Non-migratory | Mesopelagic | MR | Visual |
| Scophthalmidae | *Lepidorhombus whiffiagonis* | EF439125.1 | Non-migratory | Mesopelagic | MR | Visual |
| Scophthalmidae | *Scophthalmus rhombus* | EF439162.1 | Oceanodromous | Bathypelagic | MR | Visual |
| Scophthalmidae | *Zeugopterus norvegicus* | EU491975.1 | Non-migratory | Mesopelagic | MR | Visual |
| Scophthalmidae | *Zeugopterus punctatus* | EU638022.1 | Non-migratory | Bathypelagic | MR | Visual |
| Scophthalmidae | *Scophthalmus maximus* | EF439152.1 | Oceanodromous | Bathypelagic | BR | Visual |
| Soleidae | *Aseraggodes heemstrai* | JQ938062.1 | Non-migratory | Bathypelagic | MR | Non-visual |
| Soleidae | *Aseraggodes kobensis* | JQ938060.1 | Non-migratory | Bathypelagic | MR | Non-visual |
| Soleidae | *Brachirus annularis* | MK617127.1 | Non-migratory | Mesopelagic | MR | Non-visual |
| Soleidae | *Buglossidium luteum* | EU492030.1 | Non-migratory | Mesopelagic | MR | Non-visual |
| Soleidae | *Heteromycteris japonicus* | JQ938061.1 | Non-migratory | Bathypelagic | MR | Non-visual |
| Soleidae | *Microchirus azevia* | DQ197865.1 | Non-migratory | Mesopelagic | MR | Non-visual |
| Soleidae | *Microchirus variegatus* | EF439137.1 | Non-migratory | Mesopelagic | MR | Non-visual |
| Soleidae | *Pardachirus pavoninus* | MK617120.1 | Non-migratory | Bathypelagic | MR | Non-visual |
| Soleidae | *Pegusa cadenati* | EF456072.1 | Non-migratory | Bathypelagic | MR | Non-visual |
| Soleidae | *Pseudaesopia japonica* | JQ938063.1 | Non-migratory | Bathypelagic | MR | Non-visual |
| Soleidae | *Solea senegalensis* | DQ197904.1 | Non-migratory | Bathypelagic | MR | Non-visual |
| Soleidae | *Soleichthys heterorhinos* | JQ938064.1 | Non-migratory | Bathypelagic | MR | Non-visual |
| Soleidae | *Dagetichthys lusitanicus* | EF439470.1 | Non-migratory | Bathypelagic | BR | Non-visual |
| Soleidae | *Dicologlossa cuneata* | EF439368.1 | Non-migratory | Mesopelagic | BR | Non-visual |
| Soleidae | *Pegusa lascaris* | EF427491.1 | Non-migratory | Mesopelagic | BR | Non-visual |
| Soleidae | *Solea aegyptiaca* | JX292785.1 | Non-migratory | Bathypelagic | BR | Non-visual |
| Soleidae | *Solea solea* | DQ197905.1 | Oceanodromous | Bathypelagic | BR | Non-visual |
| Soleidae | *Synapturichthys kleinii* | EF427518.1 | Non-migratory | Mesopelagic | BR | Non-visual |
| Soleidae | *Zebrias zebra* | MK617128.1 | Non-migratory | Bathypelagic | BR | Visual |
| Soleidae | *Brachirus harmandi* | MK617105.1 | Non-migratory | Bathypelagic | FW/BR | Non-visual |
| Soleidae | *Brachirus orientalis* | JAFBJR010000150.1 | Non-migratory | Bathypelagic | FW/BR | Non-visual |

**Figure S1.** Flatfish RH1 maximum likelihood gene tree, nodes: aLRT-SH-like branch support

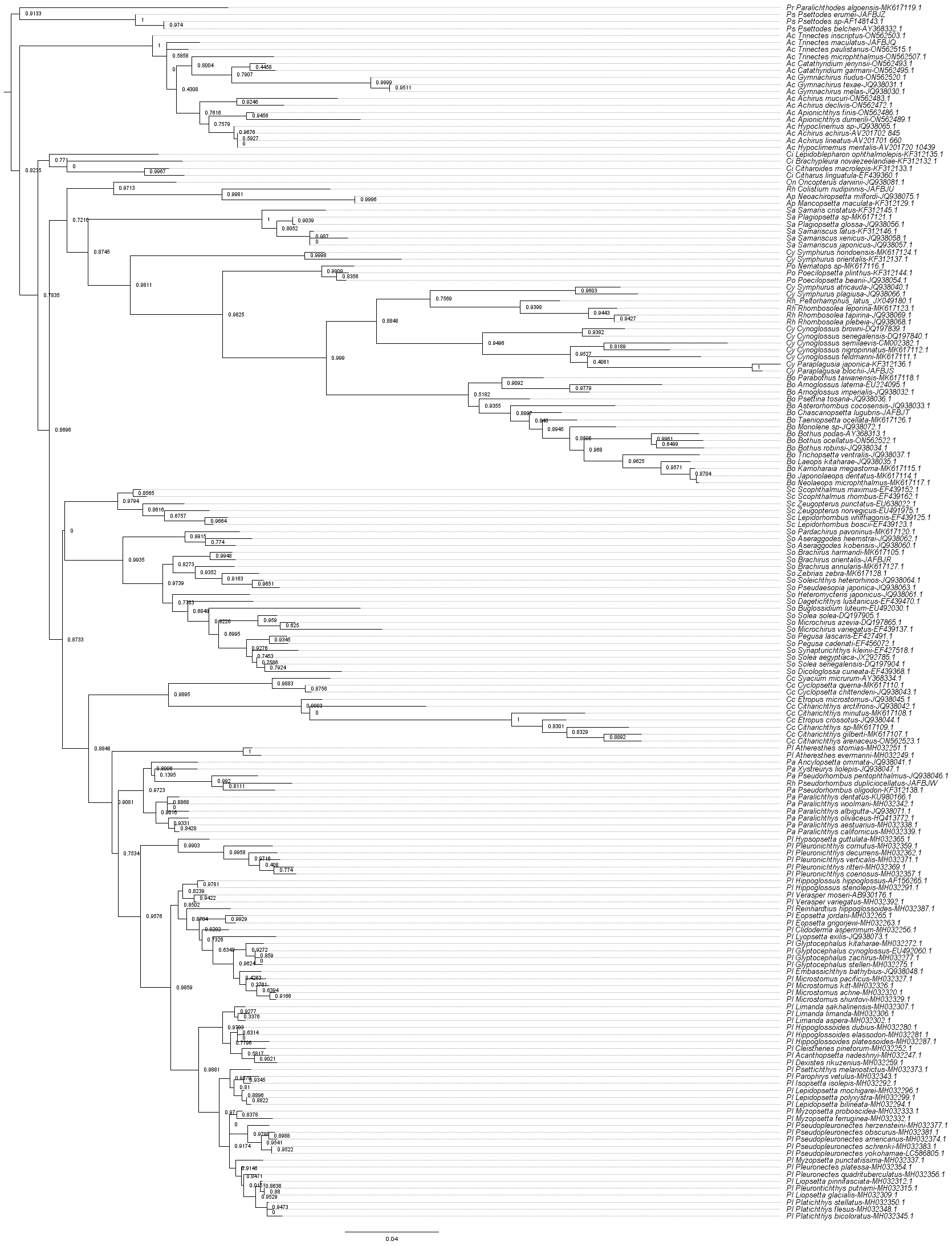
